# Energy balance implications of floral traits involved in pollinator attraction and water balance

**DOI:** 10.1101/539668

**Authors:** Adam B. Roddy

## Abstract

For most angiosperms, flowers are critical to reproduction because they increase rates of outcrossing. Flowers are highly variable in numerous traits, including size, shape, and color. Most of this variation is thought to have arisen due to selection by pollinators. Yet, non-pollinator selection is increasingly being recognized as contributing to floral trait evolution. One such non-pollinator agent of selection that often opposes pollinator selection includes the physiological and resource costs of producing and maintaining flowers. Here I (1) summarize recent studies on the macroevolution of floral physiological traits and (2) apply an energy balance model to examine how two pollination traits (flower color and flower size) can interact with hydraulic traits to influence flower physiology. These modeling results show that under certain conditions flower color variation can overwhelm the effects of floral transpiration and flower size variation on flower temperature. Additionally, the range of flower size most common in the California flora is the range in which complex, non-linear dynamics in flower energy balance occur. These results suggest that floral traits under selection by biotic agents can have large implications for flower physiology, and only some of these potentially deleterious effects can be offset by flower hydraulic traits. These complex interactions between pollination traits (flower size and color) and physiological traits (surface conductance to water vapor) suggests that a more unified framework for understanding the evolution of floral form and function would simultaneously consider the interaction between physiological traits and traits under biotic selection.

## Introduction

It is commonly assumed that an organism’s phenotype is the result of adaptation. Selection, it is thought, hones the various traits of an organism to best maximize fitness in the ecological context in which the organism thrives. The fundamental challenge facing an organism is, therefore, one of allocation: how to allocate its limited resources in order to maximize survival and reproduction. But simply observing the diversity of plant life makes obvious the fact that there is no single solution to this problem. Allocation even to reproductive structures–whose function is most immediately tied to fitness–can vary, in terms of biomass, from 1-60% among plant species (Bazzaz et al. 1987). One way to reconcile the presumption of maximizing fitness with the obvious diversity of phenotypes has been with optimality models (e.g. Parkhurst and Loucks 1972; Givnish and Vermeij 1976; Ashman and Schoen 1994). These models define cost and benefit functions, which, when solved under varying conditions, yield a variety of optimal solutions. In evolutionary models, the number of equally fit solutions increases and the maximum fitness of any one of them decreases as the number of constraints or selective agents increases (Niklas 1994). Put simply, phenotypic diversity increases as organisms must perform more tasks.

Defining the costs of plant structures for optimality models is inherently a question of physiology–what are the costs of producing and maintaining the structure? The most common currency of cost is carbon, which is a basic component of plant structures and is necessary for respiration. For photosynthetic structures, such as leaves, water is also considered a cost because the diffusion of CO_2_ from the atmosphere into the leaf requires water to evaporate. As a result, optimal leaf size and shape can be predicted for different environmental conditions based on the carbon and water costs of construction and maintenance, the carbon benefits of photosynthesis, and the various environmental factors (light, temperature, humidity) that influence the photosynthetic efficiency of leaves (Parkhurst and Loucks 1972; Givnish and Vermeij 1976). Thus, the factors that generate phenotypic diversity are those that influence the cost and benefit functions, which are typically in terms of carbon and water.

For flowers, however, benefit functions are defined by ovule fertilization and pollen dissemination (e.g. Ashman and Schoen 1994). Since the recognition and articulation of the role of flowers in plant reproduction over two centuries ago (Sprengel 1793; Vogel 1996), there has, rightfully, been a strong emphasis on characterizing the diversity of floral form and the adaptive value of various floral traits. These studies have elaborated the numerous floral traits that can be involved in pollinator interactions (Kevan and Baker 1983; Fenster et al. 2004), have detailed how different floral morphological traits are associated with different pollinators (e.g. Whittall and Hodges 2007), and have shown how certain combinations of floral characters are likely more advantageous than others (Stebbins 1951a; O’Meara et al. 2016). Yet, the costs of flowers–the resources associated with their production and maintenace (Reekie and Bazzaz 1987a,b)–have received notably less attention (Gleason 2018), despite their key role in determining fundamental traits, such as longevity (Ashman and Schoen 1994) and despite the tradeoff between resource investments in reproduction and leaf function (i.e. gas exchange and growth; Bazzaz et al. 1987; Reekie and Bazzaz 1987c; Galen et al. 1999). Furthermore, including the maintenance costs of flowers–e.g. rates of respiration–along with their static production costs can improve predictions of future reproductive effort (Ashman 1994). Thus, characterizing the adaptive accuracy of flowers for their pollination function may ignore important physiological factors that may also exert substantial influence over floral form and evolution.

Although pollinators are still considered the primary drivers of floral traits (Fenster et al. 2004), over the last few decades the role of resource availability has been increasingly acknowledged as an important factor shaping floral traits (Sapir 2017; Sapir and Ghara 2017; Caruso et al. 2018). Because non-pollinator agents of selection often oppose pollinator selection, there is often a tradeoff between these competing agents (Galen 1999; Strauss and Whittall 2006). Despite the importance of these non-pollinator agents of selection, little effort has been made to fully characterize the physiological costs of flowers and the evolution of traits underlying these costs. While studies focused on one or a few species help to elucidate physiological mechanisms (e.g. Galen et al. 1999; Chapotin et al. 2003; Roddy and Dawson 2012; Teixido and Valladares 2014a, 2015; Roddy et al. 2018), a comparative, macroevolutionary framework is critical to understanding the long-term, net impacts of variation in floral physiological traits on evolutionary dynamics and to distinguishing the developmentally possible from the selectively advantageous. Furthermore, the climate is changing faster than even the most aggressive models have predicted (Raupach et al. 2007), driving major shifts in phenology (Fitter and Fitter 2002; Parmesan and Yohe 2003) and plant investment in reproduction (Wright and Calderon 2006). Because these phenological shifts have physiological underpinnings and implications, understanding the physiology of flowers is, therefore, critical to understanding the causes and consequences of shifting flowering time on plant function and evolution.

Here I have two aims. First, I summarize recent research elucidating the physiological traits of flowers, focusing primarily on hydraulic architecture, and layout a series of hypotheses about the evolution of hydraulic structure-function relationships in flowers. That abiotic agents of selection on flowers can overwhelm the effects of biotic agents (Caruso et al. 2018) motivates a more mechanistic and more comparative understanding of the physiology of flowers. Second, I apply an energy balance modeling approach to show how traits important to pollinator interactions can have physiological implications. I focus on two traits, flower color and flower size, both of which experience selection by pollinators. My approach here is, by no means, comprehensive; rather, I aim to showcase how a physiological perspective can elucidate new insight into floral form, function, and evolution.

## Centering metabolism in macroevolutionary studies of flowers

The origin and early diversification of the flowering plants may not have presaged the grandeur of their future. The first angiosperms were most likely ruderal and adapted to living under the dominant ferns and gymnosperms (Stebbins 1951b; Feild et al. 2004). Yet the angiosperms rapidly diversified, accumulating novel traits–new reproductive structures and new anatomical features that facilitated higher rates of metabolism–that enabled them to dominate ecosystems globally and further promoted their diversity and dominance (Bond 1989; Crepet and Niklas 2009; Brodribb and Feild 2010; de Boer et al. 2012; Simonin and Roddy 2018). Traditionally, innovations in floral reproductive traits were thought to have been the primary drivers of angiosperm diversity (Sanderson and Donoghue 1994; Crepet and Niklas 2009), but more recent work has suggested that their ecological dominance–and also possibly their diversity–may be due to factors linked to metabolism (Brodribb and Feild 2010; de Boer et al. 2012; Simonin and Roddy 2018). While the earliest angiosperms were likely drought-intolerant with low physiological rates (Feild and Arens 2007; Feild et al. 2009a), subsequent lineages rapidly developed smaller, more densely packed cells that allowed unprecedented rates of primary (Simonin and Roddy 2018).

Central to this emerging perspective about early angiosperm evolution and their subsequent dominance is that the need to supply resources for primary metabolism (photosynthesis) is a main selective agent driving the structure and organization of cells and tissues. Because the diffusion of CO_2_ into the leaf mesophyll requires that stomata on the leaf surface be open, the wet internal surfaces of the leaf are exposed to a desiccating atmosphere. In this way CO_2_ uptake is mechanistically coupled to the transport of water throughout the leaf such that traits associated with water supply and loss are coordinated (Sack et al. 2003; Brodribb et al. 2005, 2007; Scoffoni et al. 2016). However, given that most flowers are heterotrophic and assimilate little carbon, selection for high rates of resource transport has likely been relaxed, potentially leading to novel combinations of anatomical and physiological traits (Olson and Pittermann 2019).

Because flowers are heterotrophic and short-lived, the higher rates of metabolism that smaller, more densely packed cells enable (Simonin and Roddy 2018) are unneccessary in flowers. Flowers have lower vein and stomatal densities and larger cells than their conspecific leaves (Lipayeva 1989; Feild et al. 2009b; Roddy et al. 2013, 2016; Zhang et al. 2018). The interspecific correlations between traits are similar within leaves and within flowers, for both anatomical and physiological traits (Roddy et al. 2016, 2019; Zhang et al. 2018), suggesting that similar scaling relationships define flower and leaf trait covariation despite divergent selection regimes acting on flowers and leaves. This divergence in flower and leaf hydraulic traits points to the importance of developmental modularity, which has allowed flowers and leaves to occupy difference regions of trait space due to their different selective constraints (Berg 1960; Roddy et al. 2013, 2019; Olson and Pittermann 2019). Furthermore, greater modularity of foliar and floral traits may have occurred among more recently derived groups. While ANA grade and magnoliid leaves have lower vein and stomatal densities than other angiosperms, their flowers have relatively high vein and stomatal densities (Figure 1; Roddy et al. 2016). The hydraulic traits of ANA grade and magnoliid flowers, therefore, seem to be more similar to those of leaves, while leaf and flower hydraulic traits of the monocots and eudicots are more different from their conspecific leaves. Though these patterns are based on relatively few species, they do suggest that metabolism has been a central agent of selection shaping flower physiology, just as it has shaped leaf physiology.

**Figure 1.**
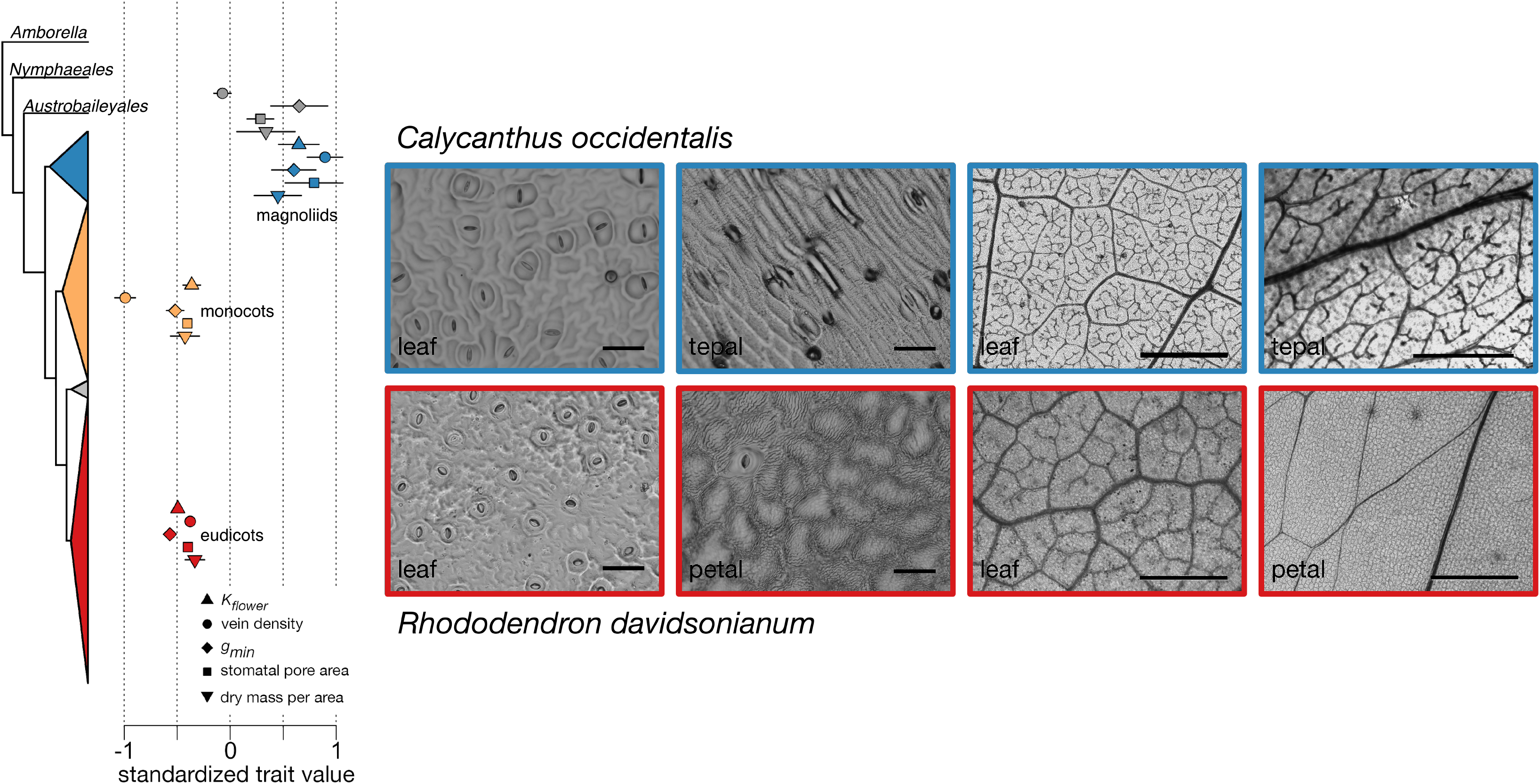
Variation in floral physiological and structural traits among extant angiosperm clades (modified from Roddy et al. (2016) with permission from the publisher). Traits are standardized and include whole-flower hydraulic conductance (*K_flower_*), vein density, minimum epidermal conductance to water vapor (*g_min_*), stomatal pore area index (a function of stomatal density and stomatal size), and corolla dry mass per projected surface area. Images show stomata (left) and veins (right) for leaves and flowers of two species, (top) *Calycanthus occidentalis* (Calycanthaceae) and (bottom) *Rhododendron davidsonianum* (Ericaceae). Outlines and colors indicate clade affinity (grey = ANA grade, blue = magnoliids, orange = monocots, red = eudicots). Scale bars for images of stomata are 100 μm and for images of veins are 1 mm.

Reducing the abundance of stomata may have been a key physiological innovation with cascading consequences on other floral traits. In leaves, stomata function to regulate water loss while CO2 diffuses into the leaf mesophyll where it is assimilated into sugars. At the same time, transpired water carries with it latent heat which helps to cool the leaf and prevent it from overheating. Volatile organic compounds (VOCs), which can function in communication, defense, and pollinator attraction, can also be released through leaf stomata (Baldwin 2010). Flower corollas, in constrast, are generally heterotrophic, and, therefore, may be relaxed from the need to transpire high rates of water to support photosynthesis. Relaxing this constraint would have allowed flowers to reduce water loss by eliminating stomata. While stomata may be important in VOC release, other structures, such as osmophores, crenulated epidermal cells, and trichomes, can also be sources of VOC release in flowers (Baldwin 2010). Eliminating stomata would have, in turn, allowed the densities of veins to be reduced (Roddy et al. 2013). This transition would have reduced both water and carbon costs of flowers and possibly would have allowed alternative modes of hydration. While *Illicium* (Schisandraceae) and magnoliid flowers are hydrated by the xylem (Feild et al. 2009a,b; Roddy et al. 2018), some have argued based on water potentials that eudicot flowers are hydrated by the phloem (Trolinder et al. 1993; Chapotin et al. 2003). Such a transition in the basic plumbing of a flower would have been a critical event in floral evolution, but unequivocal evidence for this pattern remains lacking. Further, other physiological properties, such as high hydraulic capacitance could generate similar patterns as those considered indicative of phloem-hydration (Roddy et al. 2018).

On an area basis, these transitions in both water and carbon costs seem to have occurred in the common ancestor of monocots and eudicots, although better sampling of these divergences is strongly needed (Figure 1). While reducing stomatal densities would have reduced maximum transpiration rate, stomata play a critical role in dynamically regulating water loss in leaves and flowers (Roddy et al. 2018). Without stomata, water loss from flowers may be controlled solely by epidermal and cuticular traits. The non-stomatal epidermal conductance of leaves (commonly termed the minimum epidermal conductance, *g_min_*) is increasingly being considered an important trait that may influence drought tolerance (Duursma et al. 2019), and in flowers it correlates with vein density and whole-flower hydraulic conductance (*K_flower_*), suggesting that it is key to flower water balance as well (Roddy et al. 2016). If selection has driven reduced water loss from flowers, then the loss of stomata would have drastically reduced maximum transpiration rates and any further selection on reduced water loss may have revealed epidermal and cuticular traits to selection on water balance.

The macroevolutionary patterns in floral hydraulic traits suggest that innovations in floral water balance may have been a key dimension of floral evolution among early angiosperms, but is flower water balance under contemporary selection as well? Correlations between traits and climate are often taken as strong evidence of natural selection; within species, larger flowers and floral displays occur in wetter habitats (Lambrecht 2013). Because larger flowers can more readily heat up, the disproportionate maintenance costs of large flowers can lead to shorter longevities (Teixido and Valladares 2014a, 2015), suggesting that physiological traits related to water balance interact with other traits linked to pollination success (Galen 1999, 2000; Teixido and Valladares 2013, 2014b). Insect pollinators frequently prefer warmer flowers particularly in cool climates (Kevan 1975; Dyer et al. 2006), yet flowers can overheat particularly in warm climates (Patiño and Grace 2002; Patiño et al. 2002). With morphological and physiological traits flowers must balance the need to prevent overheating and the benefit of being warmer to attract pollinators. These early results suggest that the need to maintain water balance may influence key pollination traits and affect floral form and function over both contemporary and historical timescales.

## Physiological implications of pollination traits

In cold habitats, flower shape and orientation are under selection to increase the temperature of the flower microenvironment in order to attract pollinators (Kevan 1975; Stanton and Galen 1989; Atamian et al. 2016). In many cases, then, elevated flower temperatures are beneficial, as they also can be involved in the release and dissemination of floral volatiles (Jakobsen and Olsen 1994; Sagae et al. 2008; Schiestl 2015), a component of pollination interactions that likely predates the angiosperms (Terry et al. 2007; Peñalver et al. 2012). But flower temperatures cannot exceed critical temperatures that might hinder pollination and seed development, and so there is a tradeoff between elevating temperatures to attract and reward pollinators and maintaining temperatures cool enough to prevent overheating.

Whether tall, canopy trees or small, herbaceous annuals, plants commonly place their flowers above their leaves, where air temperature is warmest and humidity is lowest. This placement can have important implications on the energy and water balance of flowers because of high solar radiation and exposure to high windspeeds. For example, in seasonally dry tropical forests, most species flower in the dry season, when rainfall and atmospheric humidity are low and air temperature and solar radiation high, creating conditions of high evaporative demand (Wright and Van Schaik 1994; Wright and Calderon 1995; Wright and Calderon 2006). In the understory, conditions are generally thought to be more stable, but even understory flowers can still overheat to temperatures that induce mortality in carpels and pollen (Patiño and Grace 2002).

One way to regulate temperature is through latent heat loss due to the evaporation of water. Leaves transpire substantial amounts of water through their stomata, but the relatively low abundance of stomata on flower petals (Lipayeva 1989; Roddy et al. 2016; Zhang et al. 2018) suggests their ability to regulate water loss and temperature is limited. Yet, stomatal densities and, thus, maximum transpiration rates, have been measured on relatively few species. Furthermore, many traits that influence energy balance (e.g. color, shape) are more variable in flowers than in leaves, suggesting that the dynamics of floral temperature may be more complicated, particularly because of the multiple selective pressures acting on flowers. One first step in understanding these dynamics would be to determine the sensitivity of flower energy balance to variation in traits linked to pollinator preference. The lower vein and stomatal densities of flowers suggest that they have lower rates of transpiration, which would hinder their ability to dissipate energy through latent heat loss. Using energy balance as a framework, I focus here on two flower traits important to pollinator interactions: flower color and flower size. In order to be seen and visited by a pollinator, a flower must stand out against the green backdrop of leaves and be distinct from other flowers in the community or from closely related species (Spaethe et al. 2001; Peter and Johnson 2008; Hopkins and Rausher 2012; Muchhala et al. 2014). Differences between pollinators in their ability to see colors can help to drive diversity in flower color, often in association with other morphological traits (Whittall and Hodges 2007), but pollinators are not the only selective agent on flower color. Similarly, flower size is commonly under positive selection from visually-oriented pollinators because larger flowers are more easily visible to pollinators, yet the physiological costs of producing large flowers can oppose pollinator selection (Galen 1999; Strauss and Whittall 2006)

### Energy balance model

To test the sensitivity of flower energy balance to variation in flower color and flower size, I modeled energy balance using the R package ‘tealeaves’ (Muir 2019). Because these models are generally applied only to leaves, there is an implicit assumption that the structure being modeled is planar, and so the temperature modeled is most applicable to a flat petal. In reality, more complex, three-dimensional shapes would cause deviation from the modeled temperatures, and these deviations could either increase or decrease temperature. For example, tubular flowers with fused petals would probably thicken the boundary layer, reduce transpiration, and elevate flower temperature (*T_flower_*) above that modeled assuming a flat flower. Other morphological traits, such as petal curvature, the presence of trichomes, and flower orientation would also influence energy balance. As a first approximation, however, I have assumed flowers to be flat and oriented perpendicular to the sky in order to estimate how variation in flower color and flower size may impact energy balance.

In all model runs, I kept constant solar radiation (660 W m^−2^, which is approximately 1500 μmol quanata m^−2^ s^−1^), air temperature (*T_air_*, 25°C), relative humidity (50%), windspeed (2 m s^−1^), and atmospheric pressure (100 kPa). Variation in each of these parameters would influence transpiration, latent heat loss, and thus, the calculated flower temperature (*T_flower_*). The modeled *T_flower_* results, therefore, reflect the relative effects of the varied parameters (color, size, surface conductance to water vapor) and not the absolute temperatures the flowers may experience.

Because such modeling is common in studies of leaves, I focus not on the effects of these environmental parameters but instead on the effects of traits important to flower function: color affects absorptivity of shortwave radiation, the total surface conductance to water vapor (*g_t_*) affects transpiration rate and latent heat loss, and flower size has complex effects on multipe parameters that affect energy balance. Two of these traits, color and size, are commonly under pollinator selection, and I examined how variation in these traits affect energy balance in combination with variation in *g_t_*. I parameterized the model with published values of *g_t_* (5, 25, 50,100 mmol m^−2^ s^−1^, which are equivalent to 0.05, 0.25, 0.5, and 1 μmol m^−2^ s^−1^ Pa^−1^ at 100 kPa atmospheric pressure) that span the range of stomatal and minimum epidermal conductances (*g_min_*) measured from a phylogenetically diverse set of species (Roddy et al. 2016, 2018). For flowers lacking stomata, the epidermal conductance is the primary contributor to *g_t_*. Further details about the energy balance calculations (e.g. characteristic dimension, absorptance) are discussed in the relevant sections below.

#### Flower color

The optical properties of flowers are influenced by a variety of factors, including surface texture (e.g. cell shape and cuticle structure; Whitney et al. 2009), internal structure of the mesophyll (van der Kooi et al. 2016), which can scatter absorbed light, and the type and distribution of pigments throughout the petal (Kay et al. 1981). These factors that influence floral color profiles can also influence flower energy balance (van der Kooi et al. 2017) and water relations. The surface texture of petals can have a significant impact on the amount of light–and, by extension, energy–that is reflected or absorbed (Gorton and Vogelmann 1996; van der Kooi et al. 2015, 2017; Vignolini et al. 2015). In general, darker pigmentation leads to higher absorptance, elevating flower temperature, but latent heat lost to transpiration can prevent overheating of the floral microenvironment (Patiño and Grace 2002).

To test for the effects of flower color in combination with variation in *g_t_* on energy balance, I modeled energy balance using the fixed parameters stated above and varied the absorptance of shortwave radiation from 1% to 90% at a fixed flower size of approximately 28 cm^2^ (equivalent to a characteristic dimension of 3 cm, assuming circular flowers). This variation in absorptance spanned almost the entire range of possible absorptances, and so to contextualize it, I measured reflectance and transmittance profiles of three *Dodecatheon* sp. (Primulaceae) flower morphs that differ in flower color (white, pink, red) in order to compare how intraspecific variation in flower color can affect absorptance while controlling for variation in other anatomical and physiological traits. I used an Ocean Optics S2000 spectrometer (Ocean Optics, Largo, Florida, USA) with sensitivity between 400 and 900 nm interfaced with an integrating sphere to measure white, pink, and red flowers. Reflectance and transmittance spectra were calculated from measurements on individual petals (approx. 1-cm^2^), having subtracted out the background spectrum with no illumination and normalized by a 99.9% reflectance standard. Absorptance profiles were calculated as the difference between unity and the sum of the reflectance and transmittance profiles. While the white flower had near uniform absorptance, transmittance, and reflectance across the visible waveband, the red and pink flowers had higher absorptance below 650 nm (Figure 2a-c). Red flowers had near complete absorptance in the 400-600 nm band. These differences in absorptance were expected to drive differences in temperature. Using a representative spectrum for incoming solar radiation measured on a sunny day, I calculated a weighted average absorptance as:

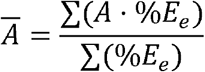

where A is the absorptance at each wavelength and %E_e_ is the intensity of that wavelength scaled to the maximum intensity of the incoming solar radiation. The weighted average absorptances for the white, pink, and red flowers were 0.29%, 26.8%, and 64.2%, respectively. The spectral profiles for these flowers are simply examples to show how color variation can affect absorptance.

**Figure 2.**
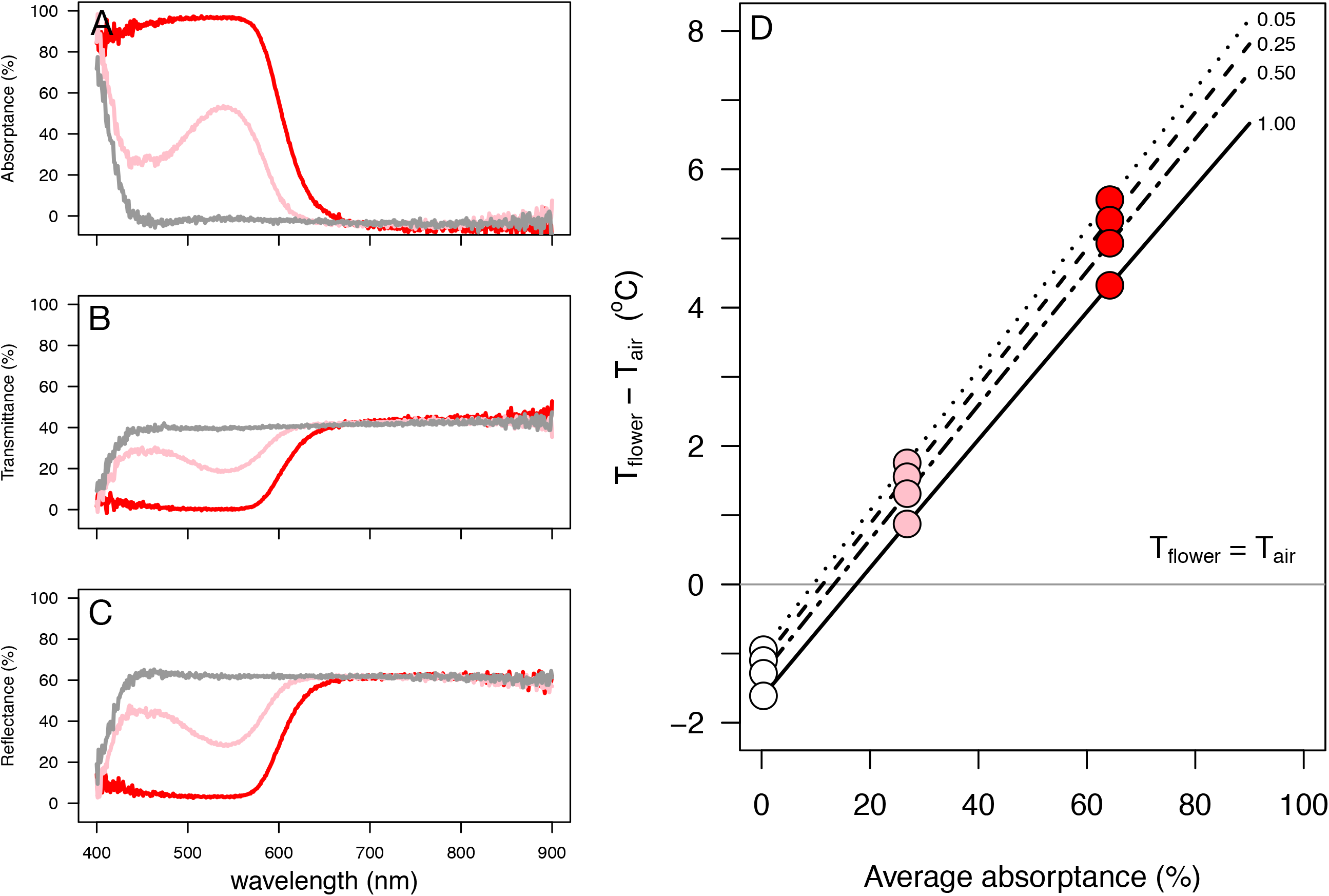
(a) Absorptance, (b) transmittance, and (c) reflectance spectra for three *Dodecatheon* sp. flowers that differ in flower color (grey lines = white flower, pink lines = pink flower, red lines = red flower). (d) The modeled differences between between flower temperature (*T_flower_*) and air temperature (*T_air_*) as a function of absorptance and the total surface conductance to water vapor (*g_t_*). Numbers indicate the value of the *g_t_* isoclines (0.05, 0.25, 0.50,1.00 μmol m^−2^ s^−1^ Pa^−1^), and colored points represent the values for the three flowers whose spectra are shown in (a-c).

While darker flowers were more absorptive, which elevated *T_flower_*, higher *g_t_* somewhat reduced *T_flower_* (Figure 2d). Darkening flowers from white to red caused *T_flower_* to increase by 6-7°C. However, varying *g_t_* twenty-fold caused *T_flower_* to change less than 1°C for white and pink flowers and approximately 1.3°C for red flowers. Therefore, the effects of flower color on energy balance can overwhelm the effects of latent heat loss due to transpiration, under the conditions modeled here. The relatively small effects of *g_t_* on *T_flower_*, relative to the effects of flower color, may explain why flowers of the monocots and eudicots have few stomata. Increasing stomatal abundance may help to prevent their flowers from overheating, but the effects would be relatively small and the benefits offset by the cost of losing water and by the potential benefit of increasing temperature to volatilizeorganic compounds. In certain habitats, however, high *g_t_* may be beneficial, such as in humid tropical environments where high humidity and low windspeeds would otherwise reduce transpiration rates and lead to elevated *T_flower_* (Patiño and Grace 2002).

#### Flower size

Despite pollinator selection for larger flowers, flowers are not universally large, and the complexity of their shape has hindered progress toward understanding the evolution of flower size. Methods for estimating flower size vary depending on the taxonomic group and the perceived importance of the sizes of different floral organs to animal pollinators. In regional floras, flower size is commonly reported as either petal length or flower diameter, yet these ways of estimating size are not necessarily physiologically meaningful. First, I tested the utility of flower size estimates available from floras by comparing measurements of the surface area of flowers to reports of flower size from the regional flora of California (Jepson eFlora, 2015). Second, I used this variation in flower size to parameterize the characteristic dimension used in the energy balalnce model. I combined variation in flower size with variation in flower color and *g_t_* to highlight the range of thermal outcomes when all three traits are allowed to covary.

First, I measured the two-dimensional, projected surface area of floral perianths on fully expanded flowers of 43 species from 29 families growing at the University of California Botanical Garden. Flowering shoots were excised, the cut shoot surfaces immediately recut underwater, and the shoots transported back to the laboratory where flowers were dissected. The calyx and corolla structures were individually placed on a flatbed scanner, and their areas measured using ImageJ (Rueden et al. 2017), totaled per flower and averaged per species. This provided two potential estimates of flower size: one in which the corolla and calyx structures were summed and one in which only the corolla structures were included. For flowers lacking a differentiated perianth, such as monocots and magnoliids, I included all tepals. For these 43 species, I extracted estimates of flower size from the Jepson eFlora (2015), which were reported as either petal length or flower diameter and often included a minimum and maximum. Because measurements of flower size were in units of area (i.e. square centimeters), reports from the flora needed to be converted into similar units for comparison. I averaged the minimum and maximum measurements reported in the flora and made two assumptions to convert these estimates into units of area: (1) that flowers are circular, and (2) that the length of a petal was equivalent to the radius of this circle. These assumptions were probably safer for actinomorphic flowers than zygomorphic flowers. I used standard major axis regression (R package “smatr”; Warton et al. 2012) to compare the slopes and intercepts in the relationships between predicted and measured flower size. Flower size estimated from the flora was a strong predictor of actual flower size, for whole flowers (slope = 0.94 [0.79, 1.11]; R^2^ = 0.63, P < 0.0001; Figure 3a) and for corollas (slope = 1.03 [0.88, 1.22]; R^2^ = 0.65, P < 0.0001; Figure 3b). For neither relationship was the slope significantly different from unity (P = 0.45 and P = 0.69, respectively), suggesting that sampling a sufficient number of species allows estimates of flower size from floras to reliably reflect physiologically meaningful estimates of flower size. This result enables mining published floras for data on flower size, a trait that is known to be important to pollination but for which there has been no consistent method for measurement.

**Figure 3.**
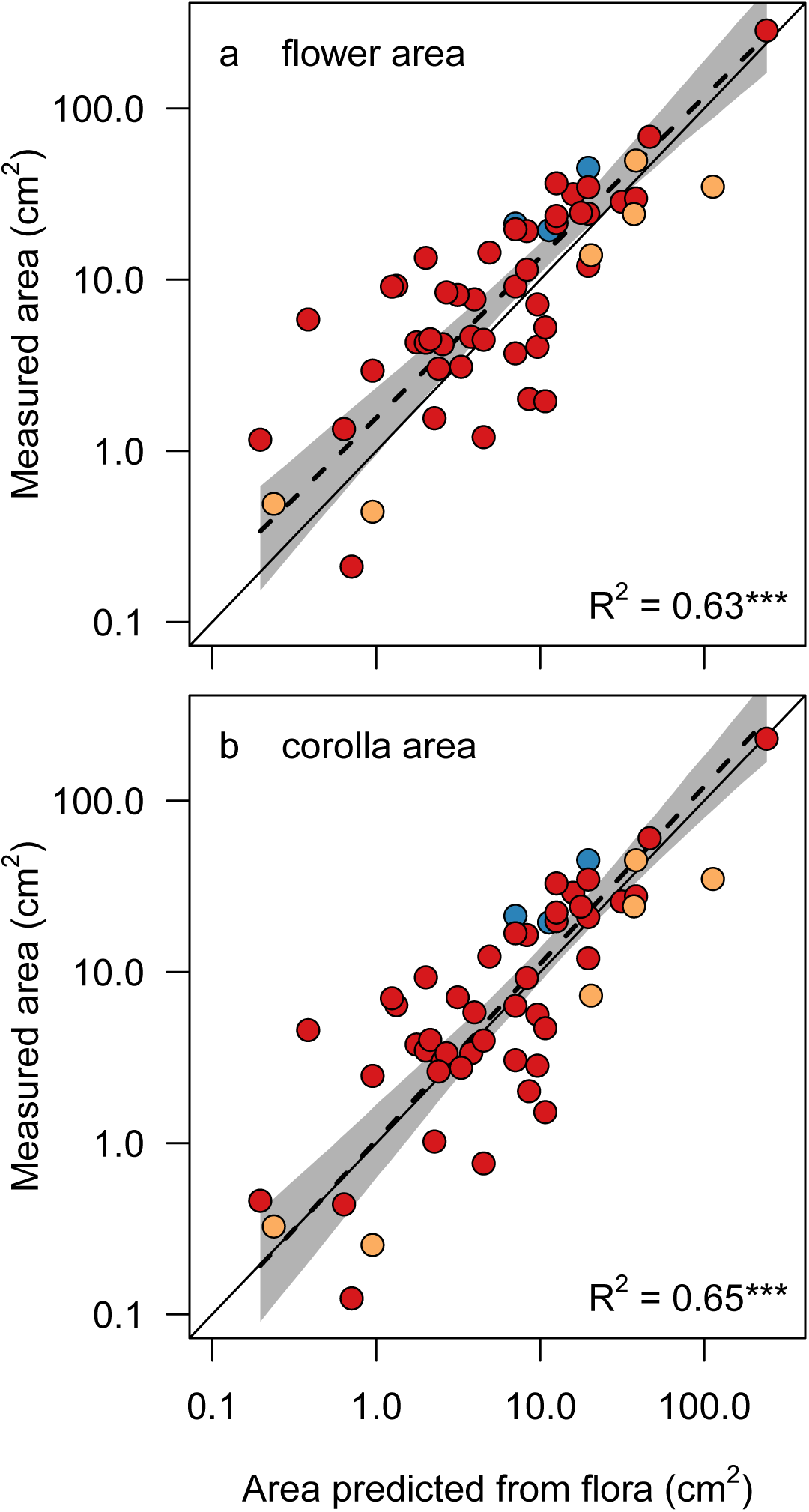
The scaling of measured flower surface area and the surface area predicted from descriptions for 43 species in the California flora. (a) The measured surface area is the total surface area of the calyx and corolla structures. (b) The measured surface area of only the corolla structures. In both figures, point colors correspond to clade membership (blue = magnoliids, orange = monocots, red = eudicots), the solid line is the 1:1 relationship, the dashed line and the grey shading represent the estimated standard major axis regression and its 95% confidence intervals.

Second, I used these measurements of flower size to to parameterize the energy balance model. Most species in the dataset had relatively small flowers, below 20 cm^2^ (Figure 4a). I varied flower size in the energy balance model by assuming that the characteristic dimension that represents size was equivalent to flower diameter. I varied this characteristic dimension from 0.5 cm to 12 cm, which represents variation in corolla surface area from approximately 0.2 cm^2^ to 113 cm^2^, the range observed among the California flora (Figure 4a). I did this for three values of absorptance, equivalent to the white, pink, and red flowers used above, and in combination with variation in *g_t_* between 0.05 and 1 μmol m^−2^ s^−1^ Pa^−1^.

**Figure 4.**
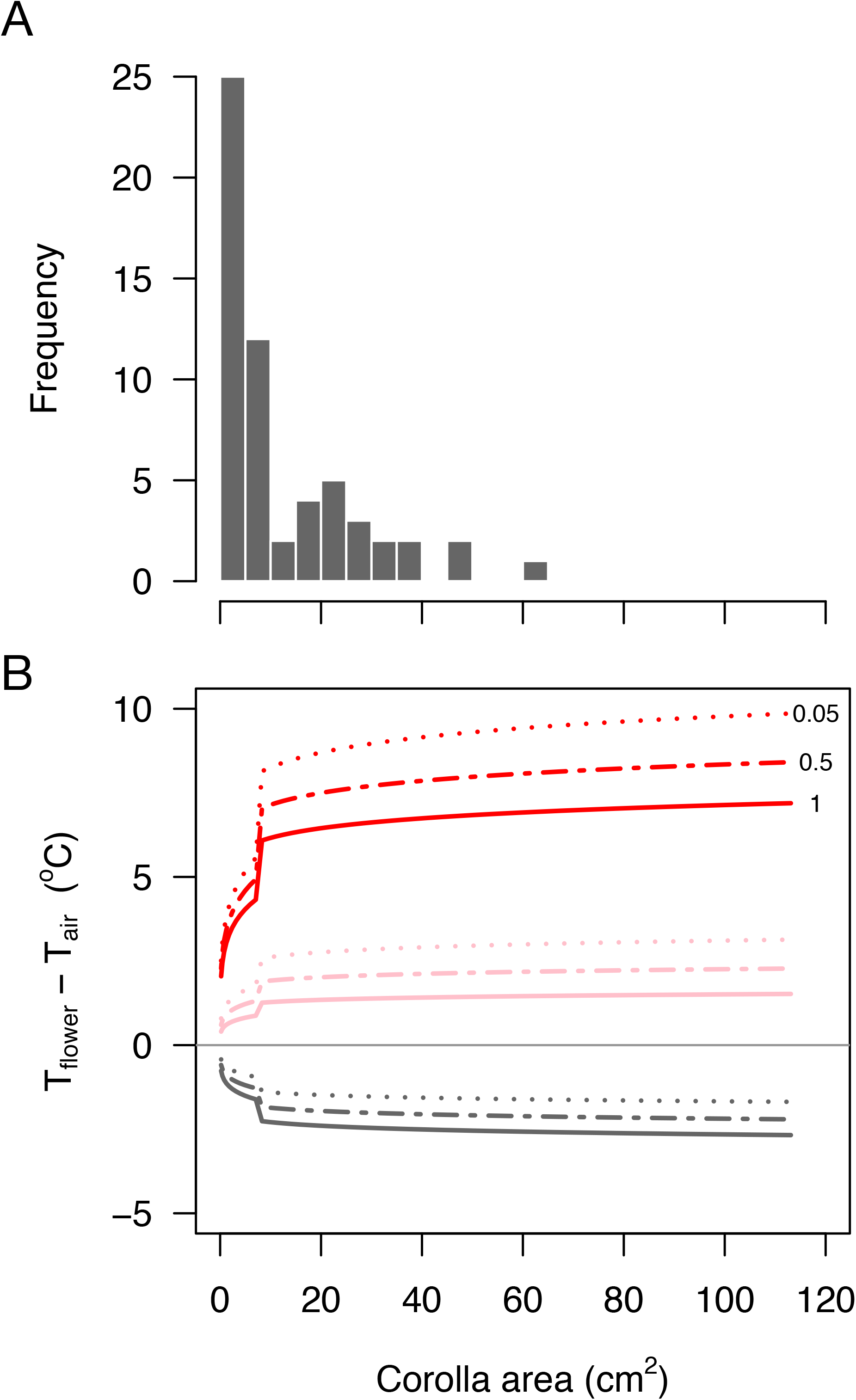
(a) Histogram of the two-dimensional projected surface area of flowers from the California flora (n = 43 species). (b) The modeled differences between flower temperature (*T_flower_*) and air temperature (*T_air_*) as a function of flower size and the total surface conductance to water vapor (*g_t_*). Numbers indicate the value of the (*g_t_*) isoclines (0.05, 0.50, 1.00 μmol m^−2^ s^−1^ Pa^−1^).

All three vairables had strong impacts on *T_flower_* (Figure 4b). Consistent with the previous results (Figure 3d), flower color had the greatest overall effect on *T_flower_*. Increasing *g_t_* universally decreased *T_flower_*, and this effect was greater for darker flowers. Above approximately 15 cm^2^ in surface area, increasing flower size had relatively little effect on *T_flower_*. Among small flowers, however, flower size variation had large and inconsistent effects on *T_flower_*. While reducing flower size among small, white flowers elevated temperatures, reducing flower size among pink and red flowers decreased *T_flower_*. Coincidentally, these large effects of small changes in flower size on *T_flower_* occurred in the range of flower sizes that are most common among the species sampled from the California flora (Figure 4a).

The morphological complexity of flowers complicates more accurate modeling of flower energy balance. For these simulations, I assumed flowers were planar, but many flowers are curved or tubular, both of which would likely reduce latent heat loss by increasing boundary layer resistance, causing an increase in *T_flower_*. In contrast, dissection (e.g. by having unfused, dissected petals) would likely reduce the boundary layer thickness and increase latent heat loss. Despite the morphological complexity among flowers, as a first approximation, this sensitivity analysis shows that nonlinear dynamics in the energy balance model may be particularly important for small-flowered species, which are predominant in certain habitats like the Mediterranean-type climate of California.

## Conclusions

Despite their ephemerality, flowers are a critical component of the plant life cycle, and their physiology is intimately tied to their performance. The physiological costs of flower production and maintenance often oppose pollinator selection and, therefore, can directly impact their form and function. Here I have used a modeling approach to examine how two traits under pollinator selection, flower color and flower size, can interact with physiological traits to impact flower energy balance. Flower color can have a large impact on flower temperature, overwhelming flower size and flower hydraulics. Variation in these traits have interactive effects, showcasing how multiple optimal phenotypes are possible. In doing so, I have shown that physiologically relevant estimates of flower size can be reliably determined from regional floras, enabling new approaches to studying flower size evolution. Overall, these results suggest that centering metabolism in our study of flowers can yield new insights into the evolution of floral form and function. For example, early evidence from macroevolutionary patterns of floral physiological traits suggests that reducing the costs of flowers would have relaxed the strength of non-pollinator selection and allowed floral traits to more rapidly and more closely track pollinator preference. These results highlight the need to more fully characterize the diverse tradeoffs between pollination traits and physiological traits. Only by more fully understanding the physiological traits of flowers will we be able to more completely understand the many dimensions of their past and future evolution in response to a changing climate.

## Acknowledgments

T.A. Kursar gave me an incredible opportunity and taught me most of the ecophysiological methods I still use; his impact is long-lasting. I thank N.M. Holbrook for the initial suggestion to work on flowers. T.E. Dawson, D.D. Ackerly, D.D. Baldocchi, P.V.A. Fine, K.A. Simonin, C.R. Brodersen, and S. Mambelli have provided valuable feedback and insight into thinking about plant form, function, and evolution. C.M. Guilliams, T. Lilittham, J. Farmer, V. Wormser, and T. Pham helped with data collection of flower size. J.H. Matthes helped with measurements of floral spectral. K.P. Tu has guided me in thinking about energy balance considerations, among other topics. I thank the organizers of the symposium from which this special issue developed, particularly Y. Sapir who gave useful suggestions on a previous draft. C.J. van der Kooi, C. D. Muir, and K. Prats also provided valuable feedback on earlier drafts.

